# A High-Resolution Probabilistic *In Vivo* Atlas of Human Subcortical Brain Nuclei

**DOI:** 10.1101/211201

**Authors:** Wolfgang M. Pauli, Amanda N. Nili, J. Michael Tyszka

**Author notes:** corresponding author: J. Michael Tyszka.

## Abstract

Recent advances in magnetic resonance imaging (MRI) methods, including data acquisition, pre-processing and analysis, have enabled research on the contributions of subcortical brain nuclei to human cognition and behavior. At the same time, these developments have led to an increasing need for a high-resolution probabilistic *in-vivo* anatomical atlas of subcortical nuclei. In order to fill this gap, we constructed high spatial resolution, three-dimensional templates, using joint high accuracy diffeomorphic registration of *T*_1_- and *T*_2_- weighted structural images from 168 typical adults between 22 and 35 years old. In these templates, many tissue boundaries are clearly visible, which would otherwise be impossible to delineate in data from individual studies. The resulting delineation provides a more accurate parcellation of subcortical nuclei than current histology-based atlases. We further created a companion library of software tools for atlas development, to offer an open and evolving resource for the creation of a crowd-sourced in-vivo probabilistic anatomical atlas of the human brain.

## Background & summary

Especially high spatial resolution magnetic resonance imaging (MRI) and improved data pre-processing, such as non-linear registration techniques [1, 2, 3, 4], now enable the targeted study of smaller human subcortical brain nuclei than was previously possible. This progress has led to an increasing need for a high-resolution probabilistic in-vivo atlas of subcortical nuclei. The need for a probabilistic reference atlas is three-fold. First, several of the relevant subcortical structures are not discernible in individual structural scans, or even when averaging structural brain scans across participants in a single study. Second, uncertainty about anatomical labels for subcortical structures hinders comparisons across studies. Third, in order to account for the small size of some subcortical nuclei, and potential spatial inaccuracies in co-registration to standard space, it is critical to maintain probabilistic atlas labels.

To address this, we improved upon an approach we recently developed to create the California Institute of Technology (CIT168) probabilistic in-vivo atlas of the human amygdala [5]. We constructed high spatial resolution, three-dimensional templates, using joint high accuracy diffeomorphic registration of T1- and T2-weighted structural images from 168 typical adults between 22 and 35 years old, released by the Human Connectome Project [6]. Subsequently, we constructed eight validation templates from 84 T1w and T2w warped image pairs, selected randomly from the full set of 168 image pairs in template space. In each validation template, three observers (WMP, AN, and JMT) labeled sub-cortical nuclei in the left hemisphere, using a previously agreed upon approach, with the Allen Brain Atlas as primary reference [7, 8]. Left hemisphere labels for each observer and validation template were mapped to the right hemisphere to generate a bilateral labeling. Each bilateral subregion label was then averaged over all 24 label volumes (eight validation templates with three observers). This resulted in a bilateral probabilistic atlas with minimal left-right observer bias.

The previous CIT168 atlas of the human amygdala has already provided a useful research tool for neuroscience studies across domains. Using a symmetric normalization (SyN) diffeomorphic image transform to map preoperative structural scans of epileptic patients onto the CIT168 atlas, it was possible to discover electrophysiological evidence for item-specific working memory activity in the human amygdala [9]. Another study, with the CIT168 atlas as reference, combined electrophysiological, lesion, and functional MRI data, and found evidence that the human amygdala processes both the degree of emotion in facial expressions and the categorical ambiguity of the emotion shown, and that these two aspects of amygdala processing can be most clearly distinguished at the level of single neurons [10]. The present extension of the CIT168 has already been used in a recent functional MRI study, which found evidence for state value prediction errors in the human substantia nigra (pars compacta), while participants solved a Markovian decision making task [11].

The present extension of the previously released CIT168 in-vivo atlas of the human amygdala [5] focuses on subcortical nuclei of reinforcement learning. Research in rodents and non-human primates has identified a network of subcortical nuclei at the core of cognition and behavior (for reviews, see [12, 13, 14]). The central tenet of this research is that a network of diverse subcortical nuclei orchestrates dopamine release in the striatum. This dopamine release implements a feedback mechanism for an individual to learn based on successes and failures in its interactions with the environment [15, 16]. More recently, advances in functional magnetic resonance imaging (fMRI) have enabled the search for analogous subcortical mechanisms in humans (for reviews, see [17] or [18]).

The present CIT168 extension includes one group of nuclei that have been found to be driving forces behind activity in the midbrain dopamine system [14, 13, 19]. These nuclei include the hypothalamus (HTH), habenular nucleus (HN), ventral pallidum (VeP), and the nucleus accumbens (NAC). Within the dopaminergic midbrain, we labeled the substantia nigra pars compacta (SNc), parabrachial nucleus (PBP), and the ventral tegmental area (VTA). We further included subcortical nuclei that are, either directly or indirectly, modulated by dopamine release, and guide behavior [20, 21, 22]. These nuclei include the caudate nucleus (Ca) and putamen (Pu), and downstream areas, including the external and internal segments of the globus pallidus (GPe and GPi, respectively), the subthalamic nucleus (STH), substantia nigra pars reticulata (SNr), and the thalamus. In addition, the atlas includes labels of subcortical structures that act as landmarks. This last group includes white matter tracts running in-between the structures of interest, and nuclei that may be misidentified as one of the above nuclei of interest. See Figure 1 and the Method section for an overview and detailed discussion of labeled nuclei.

**Figure 1:**
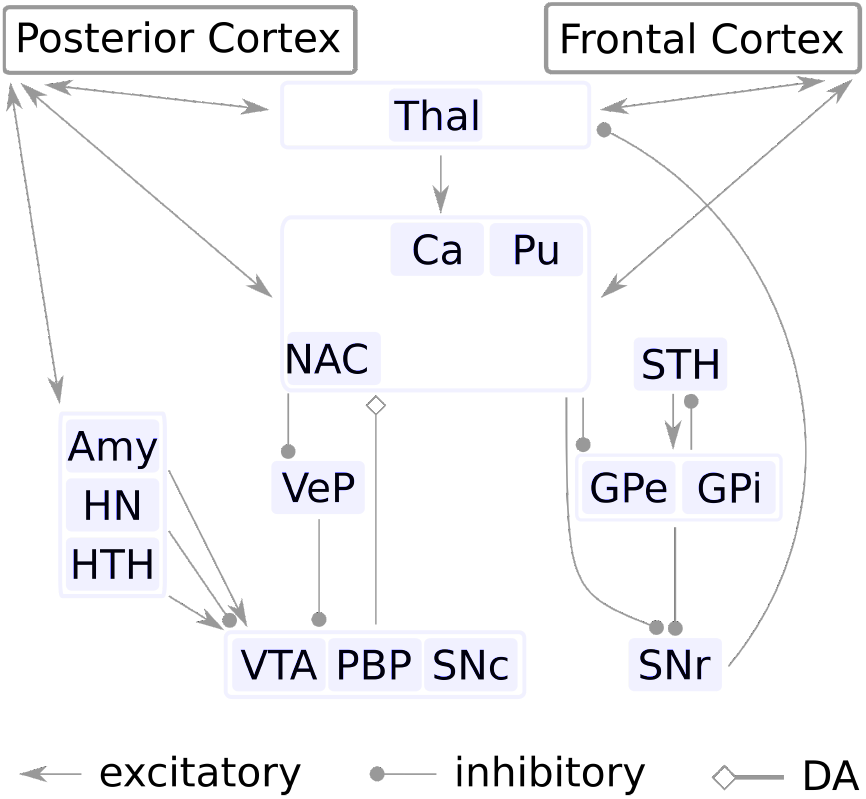
Simplified schematic illustration of subcortical nuclei and their interactions during reward learning and decision making (see main text for discussion). See Table 1 for acronyms.

In summary, we strove to create a probabilistic in-vivo reference atlas for subcortical nuclei involved in reward learning and decision making, but not included in existing in vivo atlases [23, 24, 25]. The current delineation provides a finer parcellation of subcortical nuclei, with more accurate external boundary definition than current histology-based atlases, when used in conjunction with high accuracy registration methods, such as diffeomorphic warping. These templates and delineation are intended to be an open and evolving resource for future functional and structural imaging studies. In particular, we have created a library of Python tools, which allow collaborative improvements of the present atlas, both in terms of increasing the accuracy of included anatomical labels, but also to include additional brain structures.

## Methods

### Source data

All structural images were obtained from the Human Connectome Project (HCP) S500 subject release, August 2014 [6], which includes 526 MRI datasets from individual adult human subjects, including 700 *µm* isotropic *T*_1_-weighted (T1w) and *T*_2_-weighted (T2w) whole brain images. The only inclusion criterion applied was that the T1w and T2w structural images were constructed by simple averaging of two single-average images following 6-parameter rigid body alignment [26], which reduced the available sample to 208 subjects. Age and sex unbiasing was performed by balancing the number of male and female subjects at each integer age between 22 and 35 years old (inclusive), resulting in a final sample of 168 individuals (84 males and 84 females, mean ± sd age in both groups = 28.9 ± 3.6 years). All structural images provided by the HCP S500 data release were gradient non-linearity and radiofrequency (RF) bias corrected, rigid-body AC-PC aligned, and readout distortion corrected with accurate co-registration of the individual T2w to T1w imaging spaces [26]. Sinc and spline-based interpolation was used throughout by the HCP preprocessing pipeline to minimize cumulative smoothing from repeated resampling. However, some residual blurring and spatial noise correlation are inevitable in the final individual T1w and T2w images used to construct the templates below.

### Group template construction

An unbiased template was constructed using diffeomorphic registration with a joint cost function over both T1w and T2w high-resolution 3D images from all 168 retained subjects. All registrations were performed using the bivariate symmetric normalization (SyN) algorithm implemented by the Advanced Normalization Toolbox (ANTs) [4]. Initial unbiased seed templates were constructed for T1w and T2w volumes by simple averaging across all subjects as all volumes were already rigid-body AC-PC aligned (i.e. without linear scaling) by the minimal HCP structural preprocessing pipeline.

The initial unbiased bivariate template was refined iteratively by the concatenation of affine and diffeomorphic registrations of individual T1w and T2w structural images to their respective templates generated by the previous iteration. A single diffeomorphic mapping was optimized for each individual brain using a joint cross-correlation similarity metric with equal weighting to the T1w and T2w images [27]. Only a single diffeomorphism is required as the individual T1w and T2w images were accurately coregistered during HCP preprocessing [26]. The velocity field of the diffeomorphic transform was regularized using a local Gaussian-weighted kernel with *σ* = 3.0 voxels to avoid overfitting the warp field to image noise [28]. It should be noted that the ANTs template construction pipeline includes normalization to the whole volume mean intensity and a Laplacian edge enhancement filter by default. The purpose of the edge enhancement is to compensate for blurring induced by intensity averaging alone, and serves the same purpose as the blurring inversion proposed by Avants et al. for the symmetric group normalization (SyGN) algorithm in [29]. Both approaches result in edge-enhanced templates that emphasize both the shape and appearance of anatomical structures. The template refinement was terminated after four iterations, resulting in AC-PC aligned T1w and T2w unbiased templates with 700 *μm* isotropic spatial resolution. These templates are subsequently referred to as the CIT168 templates.

### Probabilistic atlas construction

Some of the included subcortical nuclei are difficult to delineate in individual structural images from the HCP dataset. In contrast, most nuclei become sufficiently well-defined, either directly, or because of well-defined surrounding subcortical structures. This enables manual labeling in unbiased bivariate templates generated from approximately 80 or more registered individual structural images [5]. The final iteration of the joint template construction results in 168 T1w and T2w image pairs warped from individual spaces to the template space, which are then averaged to generate the final T1w and T2w templates. Consequently, we constructed validation templates from 84 T1w and T2w warped image pairs, selected randomly from the full set of 168 image pairs in template space. The unselected 84 image pairs were used to construct complementary T1w and T2w templates, which were also used for labeling validation. This process was repeated with new random samples to generate eight (four pairs of complementary) T1w and T2w templates for manual labeling by three experienced observers. The number of validation templates was considered to be a reasonable balance between total labeling time (typically 4 to 8 hours per template per observer) and the need for intra-rater validation.

All three observers (AN, JMT and WMP) labeled subcortical nuclei in the right hemisphere in each of the eight validation templates using an agreed upon, ordered approach, with the Allen Brain Atlas as primary reference. The joint unbiased T1w and T2w templates were viewed simultaneously in ITK-SNAP (version 3.6.0) [30] using a yoked 3D cursor allowing tissue volumes to be defined by referencing both contrasts. Labeling was performed in the right hemisphere only. Because each of the eight validation templates were constructed in the master CIT168 template space (see above), the probabilistic labels for each division were constructed by simple averaging over all manually labeled volumes.

Right-hemispheric labels for each observer and validation template were mapped to the left hemisphere using the following approach. Each T1w and T2w validation template was reflected about the mid-sagittal plane and warped to its unreflected version using a joint cost function affine and diffeomorphic transform. The combination of a reflection, followed by an affine and then a diffeomorphic transform (reflection warp) results in an anatomically constrained, high accuracy mapping of points in the right hemisphere to their homotopic counterparts in the left hemisphere. The reflection warp is then applied to the observer labels with nearest neighbor interpolation to generate a bilateral labeling. Each bilateral subregion label was then averaged over all 24 label volumes (eight validation templates with three observers). This resulted in a bilateral probabilistic atlas with minimal left-right observer bias (any unique observer variations in the left hemisphere would be duplicated in the right hemisphere). Labeling only one hemisphere is not without precedence in the subcortical atlas literature [31, 32]. The accuracy of reflection warping is addressed in previous work [5].

### Region delineation

The primary reference for regional delineation was the Allen Institute Adult 34 year old human atlas available at http://brain-map.org. Tissue boundaries were characterized as either explicit, with clear contrast between neighboring regions, or implicit, where low tissue contrast forces the observer to use surrounding landmarks to estimate the boundary location when labeling (see Figure 2).

The present extension of the CIT168 atlas consists of 16 gray matter regions labeled by three observers in eight templates. Template construction is described in detail in [5]. The regions and the approach to delineation for each is detailed below (see also Figure 3). The following paragraphs provide a discussion of labeling criteria, as well as the motivation for including them in the current release. For the majority of subnuclei, this motivation is based on the established role of each of these nuclei in reinforcement learning in non-human animals.

#### Caudate Nucleus (Ca)

The Ca receives strong dopaminergic innervation from the ventral tier of the SNc [33]. It receives synaptic input from prefrontal areas, and also exerts a modulatory influence over prefrontal areas, via the output nuclei (see below) of the basal ganglia [34, 35]. The Ca is involved in various goal-directed behavior and cognitive functions [36, 37, 38]. The Ca is well-defined in both T1w and T2w templates.

#### Putamen (Pu)

The Pu receives strong dopaminergic innervation from the ventral tier of the lateral SNc [33]. The Pu is critical for the execution of motor behavior [39, 40]. The Pu is well-defined in both T1w and T2w templates. We include the ventral putamen in the main Pu label, rather than as a separate anatomical label.

#### Nucleus Accumbens (NAC)

The NAC is bidirectionally interconnected with the VTA and the dorsal tier of the SNc [33]. It thus is both the target of strong dopaminergic innervation, and also modulates dopamine release in dorsal striatal areas [33]. The boundaries between the nucleus accumbens and the caudate and putamen respectively are indistinct, but not entirely invisible.

**Figure 2:**
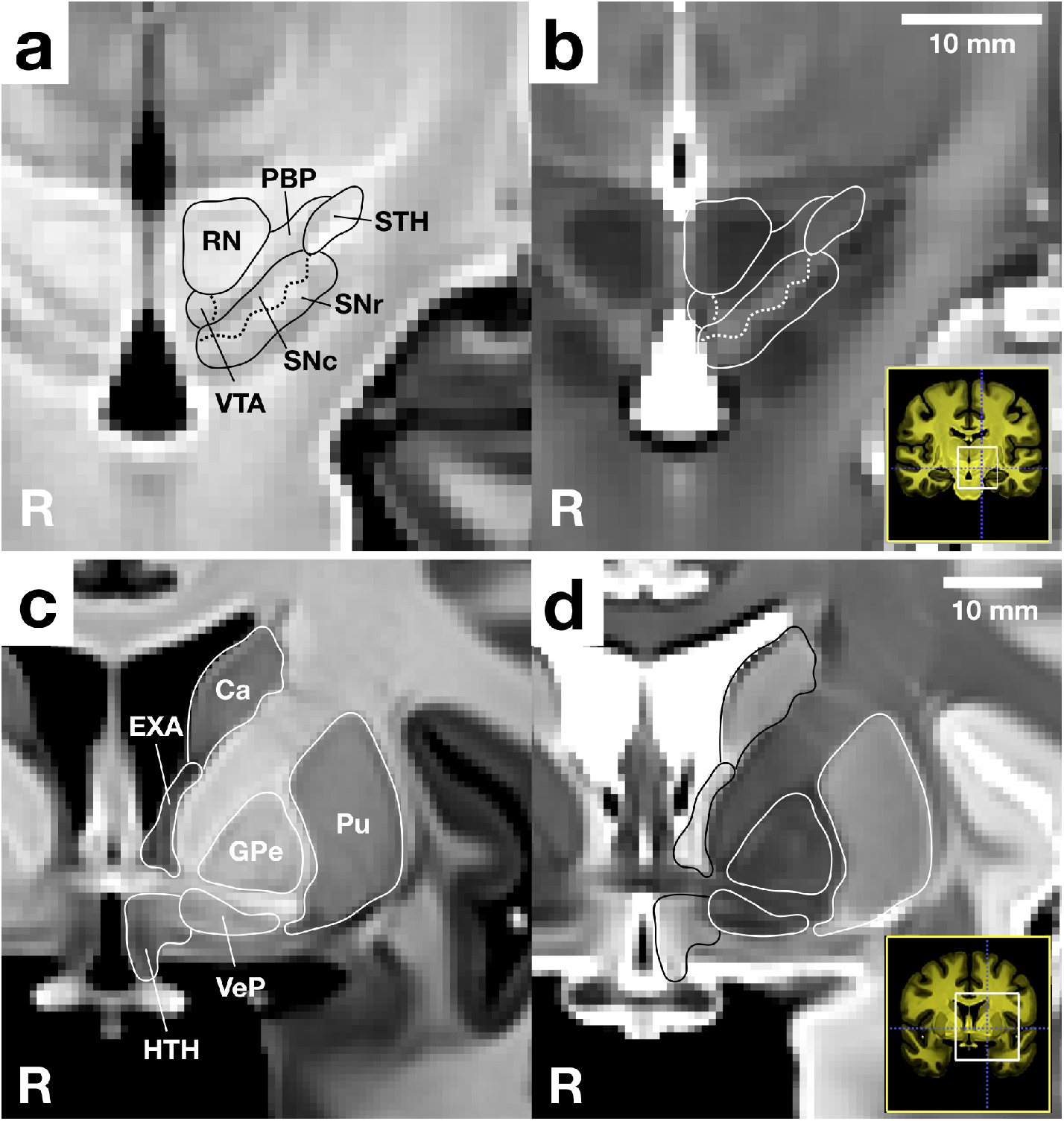
Example explicit (solid lines) and implicit (dashed lines) boundaries between the red nucleus (RN), parabrachial pigmented nucleus (PBP), substantia nigra (SNc and SNr) and subthalamic nucleus overlayed on the CIT168 T1w and T2w templates [5]. The isotropic voxel size is 700 *μm*. See Table 1 for definitions of label acronyms.

**Figure 3:**
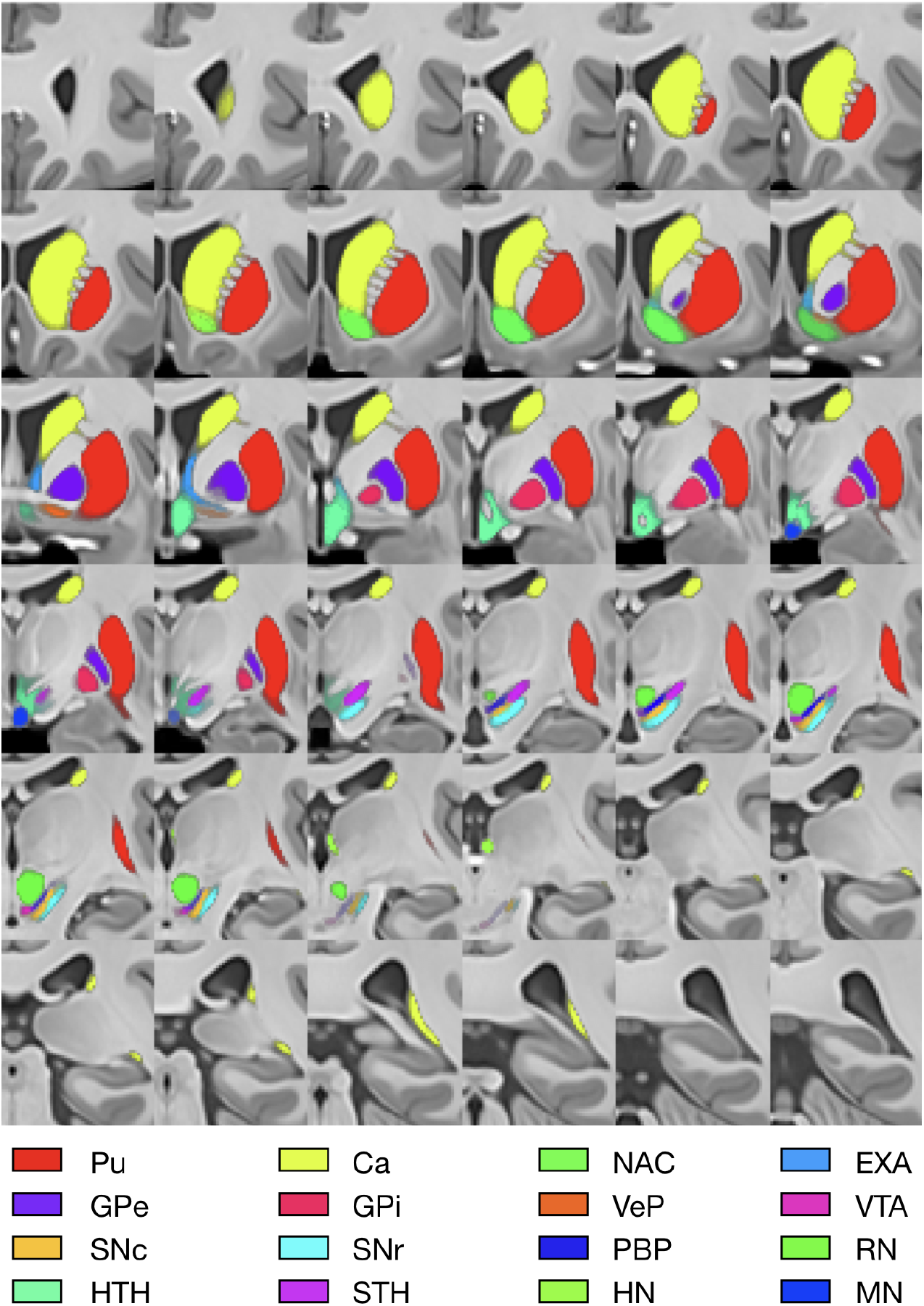
Coronal montage of probabilistic subcortical nuclei that were added in this release of the CIT168 atlas, overlaid on the T1w template. See Table 1 for definitions of label acronyms.

#### Ventral Tegmental Area (VTA)

The VTA is rich in dopamine neurons, and triggers dopamine release in the ventral striatum [33, 41]. The VTA is the target of projections from the shell of the nucleus accumbens [33], as well as from the central nucleus of the amygdala [42]. The VTA is ventral to the RN and ventromedial to the PBP. The boundary with the RN is well defined (explicit boundary), but the transition from PBP to VTA cannot be resolved in either T1w and T2w templates and must be inferred (implicit boundary).

#### Parabrachial Pigmented Nucleus (PBP)

Similar to the VTA, the PBP is rich in dopamine neurons [43], and also projects mainly to the ventral striatum [41]. The PBP is visible as a dark band in the T1w templates, dorsomedial to the SNc. The PBP has a high contrast boundary with the red nucleus dorsomedially and a lower contrast boundary with the SNC ventrolaterally.

#### Substantia Nigra, pars compacta (SNc)

The SNc is lateral to the VTA, between the SNr and the PBP. Similar to the VTA, it contains predominantly dopaminergic neurons, and triggers dopamine release in the striatum [33]. Previous work has shown an important distinction between a dorsal and a ventral tier within the SNc [33]. However, it is not possible to distinguish these tiers in vivo in individual structural images by MRI. The SNc is visible in the T1w templates as a semi-continguous, irregularly-shaped, light band.

#### Substantia Nigra, pars reticulated (SNr)

Within the frontal cortical - basal ganglia circuitry [34], the SNr receives inhibitory projections from the striatum [44]. Neurons in the SNr are tonically active and exert tonic inhibition of the thalamus [44]. Therefore, activation of striatal projection neurons cause a disinhibition of the thalamus, which is thought to produce a gating function [45, 46, 47] for working memory and motor programs in cortical areas [48]. The SNr is visible as a darker band in the T1w templates, ventral to the SNc.

#### Globus Pallidus (GP)

The internal and external segments of the globus pallidus are considered the output nuclei of the basal ganglia [39, 49]. Both the internal (GPi) and external (GPe) segments of the globus pallidus have well-defined margins in both the T1w and T2w templates.

#### Subthalamic Nucleus (STH)

The tonic activation of neurons of the SNr is at least partially attributable to excitatory input from the STH [39, 50]. The STH is well-defined in both the T1w and T2w templates. Because the ventromedial boundary of the STN with SNr and PBP is indistinct in the T2w templates, segmentation of the STH relied on combined evaluation of the T1w and T2w contrast, as this boundary is clearly visible in the T1w contrast.

#### Hypothalamus (HTH)

The HTH is involved in a diverse set of functions that are beyond the scope of this report. Briefly, it is involved in hormone release, control of food intake, fear processing, and sexual behavior. Within the scope of reinforcement learning, the lateral HTH is thought to mediate the delivery of rewards [51], and to cause dopamine release in correspondence with a reward prediction error [52]. The hypothalamus is internally heterogeneous in both T1w and T2w templates and is best defined in relation to external landmarks including the anterior commissure, mammillary nucleus, extended amygdala, sublenticular fascicle and thalamus.

#### Habenular Nuclei (HN)

The HN have been found to exert a modulatory influence over the dopaminergic midbrain [53]. The lateral HN has been hypothesized to play an role during aversive learning [54]. In the context of reward learning, it has been shown to be involved in predicting the exact timing of reward delivery [55, 54, 56]. The habenular nucleus, or habenula, is well-defined in both T1w and T2w templates.

### Neighboring structures

Several neighboring structures were also included in the probabilistic atlas, so that they could be used as landmarks during atlas construction, but also for researchers or students who are using the atlas for their work. The red nucleus (RN) is well defined in the T2w template but almost invisible in the T1w templates. It is an important landmark for the PBP, SNr, SNc and VTA labels. The mammillary nucleus (MN) is well defined in both T1w and T2w templates. The extended amygdala (EXA) consists of the bed nuclei of the stria terminalis (BNST) and the sublenticular extended amygdala (SLEA). The interstitial nucleus of the posterior limb of the anterior commissure, typically considered part of the extended amygdala, is poorly resolved by MRI and is excluded accordingly. The anterior commissure (AC) is well-defined in the T1w template and separates GPe/i from VeP. We further delineated the fornix (FX), as it provides a useful reference structure for the hypothalamus (HTH), which is otherwise somewhat indistinct. Similarly, we also labeled the optical tract (OT). The ventral pallidum (VeP) is internally heterogenous in both T1w and T2w templates, partially due to its relatively small size, and edge effects with surrounding tissue introduced during midspace template construction (see Methods “Group template construction”). However, it is not entirely indistinct, and can be well localized because of surrounding nuclei. Rostrally, it borders with the nucleus accumbens. Ventrally, it borders with the hypothalamus and substantia innominata (nucleus basalis of meynert). Dorsally, it borders with the anterior commissure.

**Table 1:**
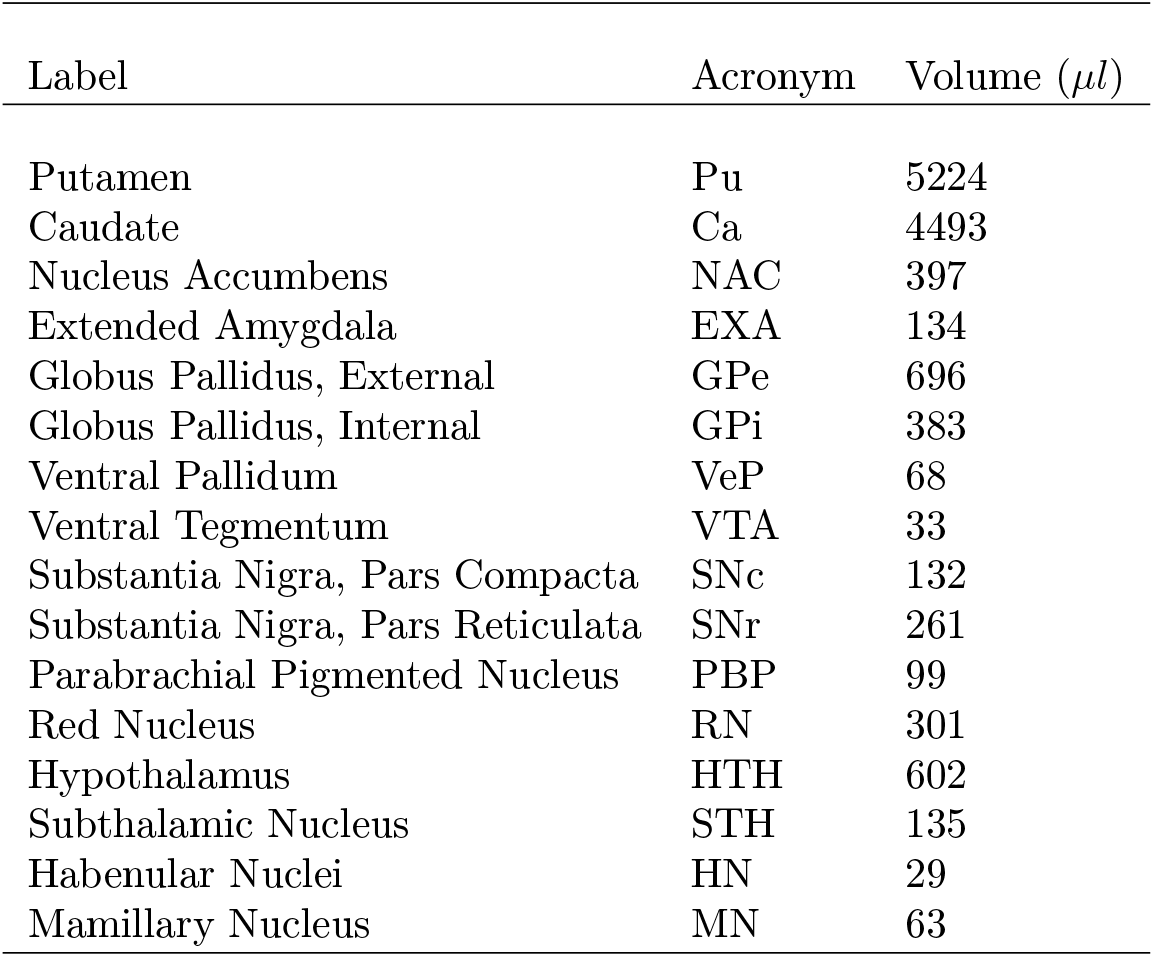
Volumes of subcortical nuclei: We derived the volume of each subcortical nucleus of the left hemisphere in microliters by spatial integration of label probabilities.

### Volumes of subcortical nuclei

In order to assess the size of the different subcortical nuclei, we estimate the volume of each of the subcortical nuclei by spatial integration of the their voxelwise probabilities over the entire volume (Table 1).

### Cumulative probability distribution of each atlas label

The construction of a probabilistic rather than deterministic atlas helps encode observer uncertainty in a natural and well-established way. The cumulative density distribution of probabilistic atlas labels provides a representation of this uncertainty (Figure 4).

**Figure 4:**
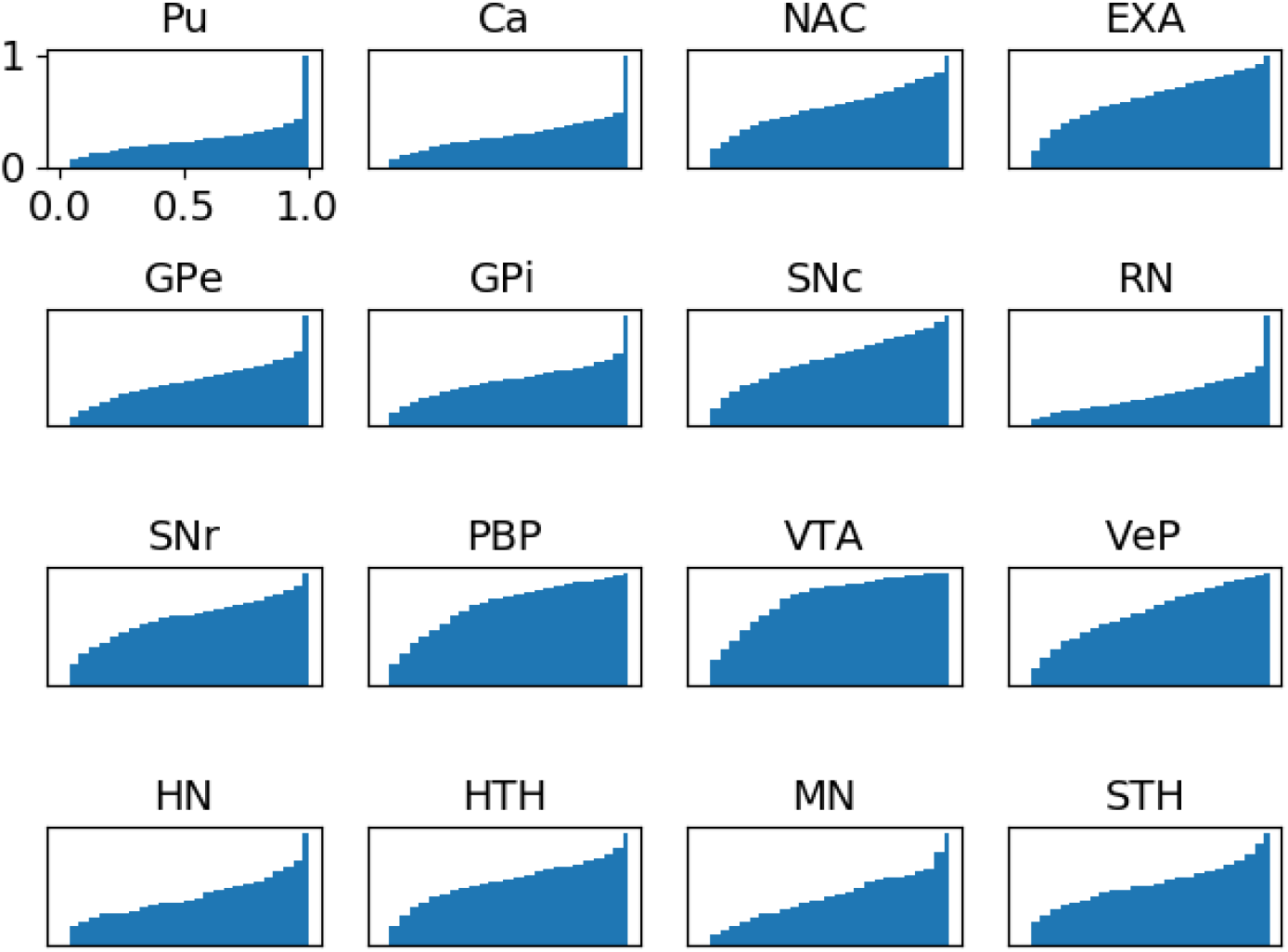
Cumulative probability distributions: To provide a representation of the uncertainty around each subcortical nucleus label, we calculated the cumulative probability distribution across the membership probability for each voxel associated with a subcortical label. Pu and Ca are examples of labels with a high degree of both inter- and intra-rater similarity, while the more convex cumulative distributions observed for PBP and VTA reflect increased inter-rater variance as the label volume decreases and the tissue boundaries become less reliably defined.

### Code availability

Tools and utilities for generating probabilistic atlases from segmented templates, and for evaluating inter- and intra-rater reliability, are maintained in a publicly available code repository https://github.com/jmtyszka/atlaskit.

## Data Records

The probabilistic atlas, including anatomical images, T1w and T2w templates, as well as segmentations of labels are available at the Open Science Framework (OSF) http://osf.io/jkzwp. We invite contributions by other researchers, in terms of alternative opinions on labeling included subcortical nuclei, as well as inclusion of additional subcortical nuclei.

## Technical Validation

Intra- and inter-rater labeling reliability between equivalent labels was assessed using two similarity measures: 1. the Dice coefficient, D, (also known as the Sörensen index), and 2. the directed or forward Hausdorff distance, H. The Dice coefficient, D, is defined as a ratio of the intersection volume of two labels to the mean volume of the two labels, in the range [0,1] [57]. To calculate H between two labeled regions, A and B, we first determine for each voxel in A the minimum Euclidean distance to any voxel in B, and then determine the maximum of all these minimum distances. This definition of H is sensitive to the ordering of A and B, and is therefore typically referred to as the directed Hausdorff distance [58]. H has identical units to the voxel dimensions, and is a measure of proximity between two regions which takes account of shape and orientation. It finds frequent application in machine vision to locate a template object within a scene [59]. Inter- and intra-rater Dice and Hausdorff metrics are conveniently viewed as matrices for each label. Example inter-rater Dice and Hausdorff metrics for the first template are shown in Figures 5 and 6 intrarater labeling reliability for observer AN is shown in Figure 7. Full similarity metrics for all observers and templates are provided in the OSF repository (see Data Citations below). Summary statistics for inter- and intra-rater similarity metrics are presented in Table 2 and 3.

It should be noted that the Dice coefficient is sensitive to the average volume of the two regions being compared. As the average volume decreases, small errors in overlap begin to dominate, until in the extreme of a single voxel label, an overlap error of only one voxel results in a Dice coefficient of zero. We therefore do not expect Dice coefficients for small volume labels to approach those typically encountered for large volume labels, such as brain masks, which routinely exceed 0.95 [60]. The Hausdorff distance is sensitive to small outlier regions present in one label only, so that even a single voxel at a distance from the main label region can skew the final metric if it is not present in the compared label volume. Taken together, the two metrics provide complementary information about label shape, positioning and overlap similarities within and between observers and between atlases.

**Table 2:**
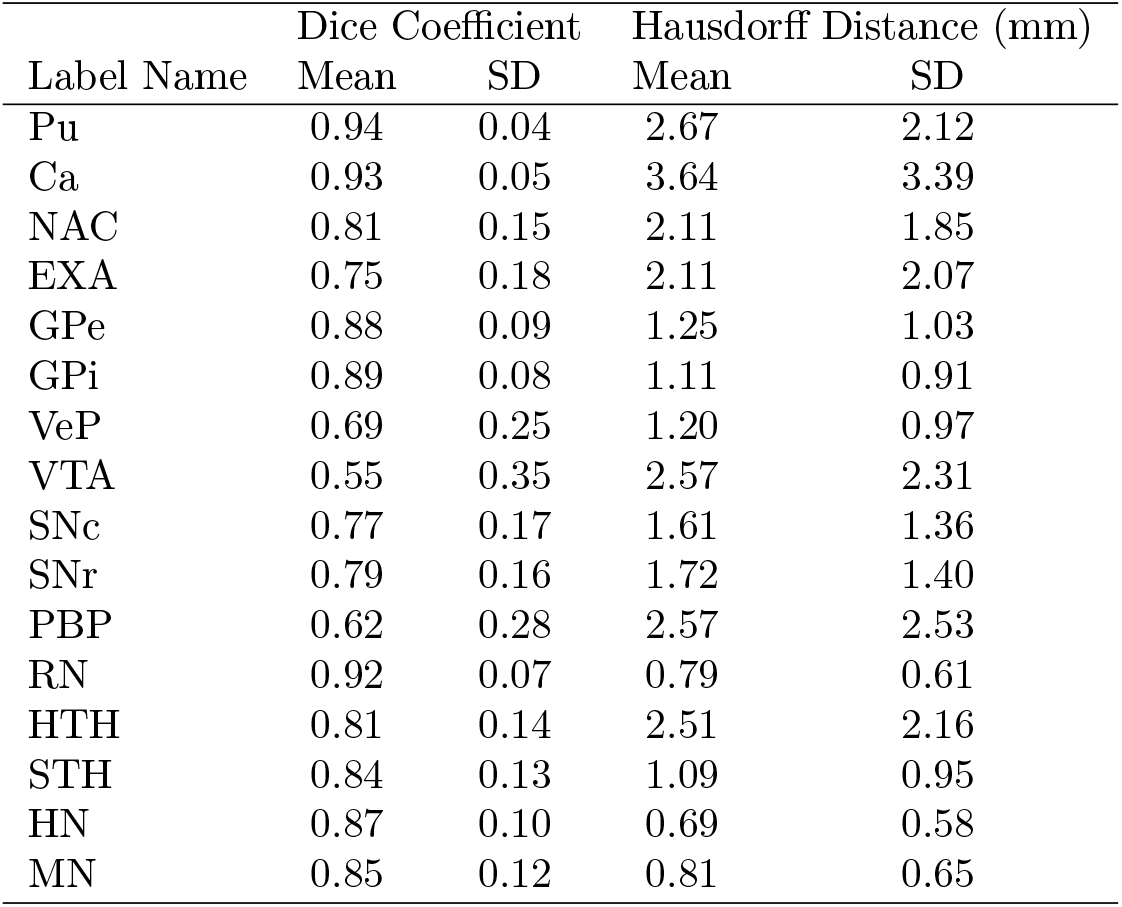
Inter-rater Similarities: Summary statistics for inter-rater Hausdorff distances and Dice coefficients averaged over all observers and templates. See Table 1 for acronyms.

## Usage Notes

We strongly recommend using the probabilistic labels as weights for calculations, rather than converting to maximum-likelihood or thresholded deterministic labels. One of the main reasons for this recommendation is that the SNc/SNr boundary is highly inter-digitated in individual brains, a feature that is entirely eliminated by averaging in the T1w and T2w templates. Consequently, deterministic labels (binary or integer valued) give a false impression of precision for the SNr/SNc boundary when mapped back to individual brains, and should be avoided where possible.

We ask researchers who would like to contribute to this evolving resource to visit the project on the open science framework (OSF) http://osf.io/jkzwp. Briefly, contributers will be asked (1) to download the validation templates, (2) to label their brain region of interest in their preferred tool (e.g. ITK-SNAP), to then either (3a) upload the labeling files to the OSF project page, or to (3b) send them to the corresponding author.

**Figure 5:**
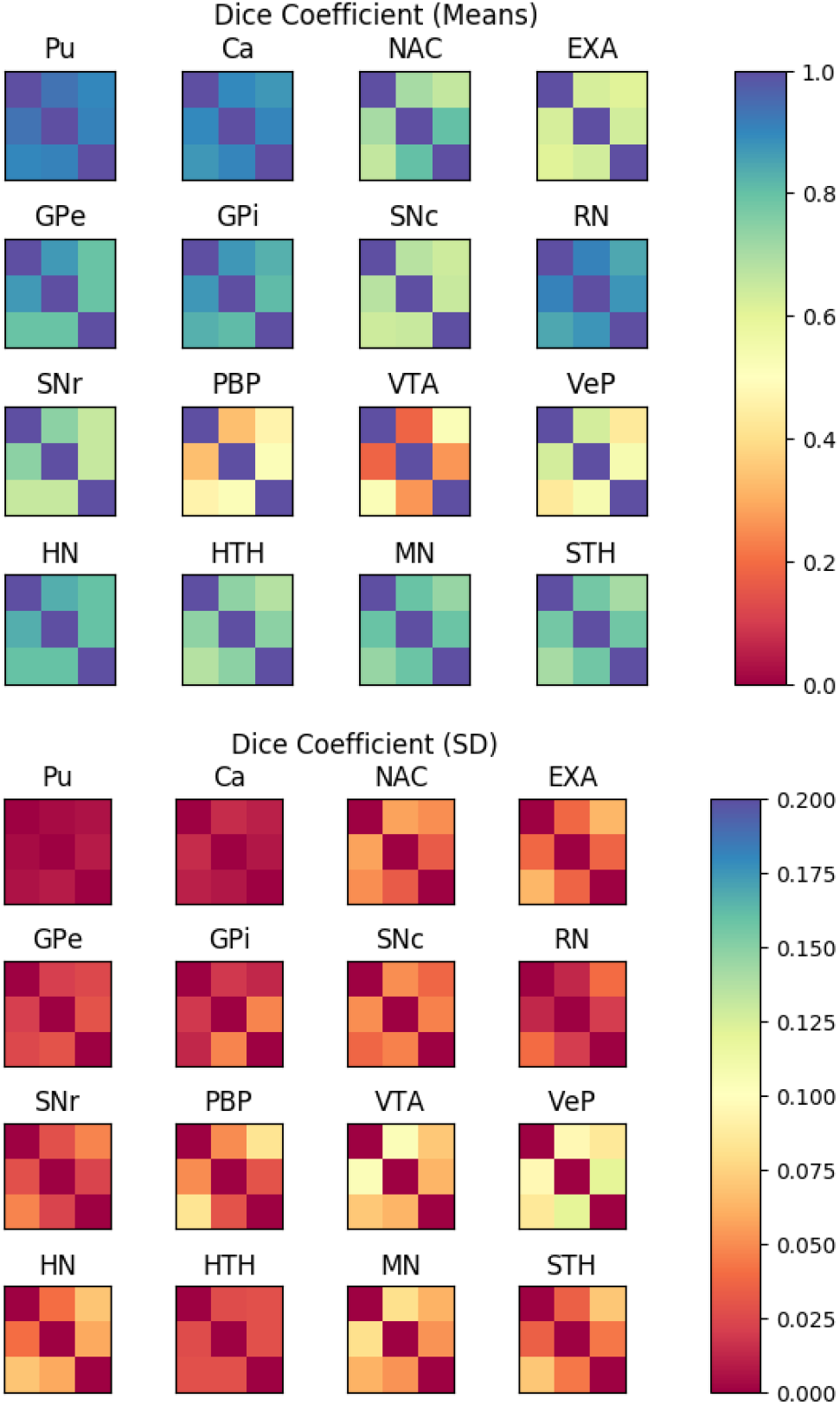
Inter-rater Dice Coefficients: Means (top) and and standard deviations (SD; bottom) of Dice similarity coefficients among raters of the 16 main subcortical nuclei included in this atlas. Each row and column correspond to one rater (AN, JMT, and WMP). Each rater labeled subcortical nuclei in each of the eight validation templates. Means and standard deviations were calculated by aggregating Dice similarity coefficients across templates. Underlying inter-rater Dice coefficients for the different validation templates can be found in the OSF repository (see Data Citations). See Table 1 for acronyms of subcortical nuclei.

**Figure 6:**
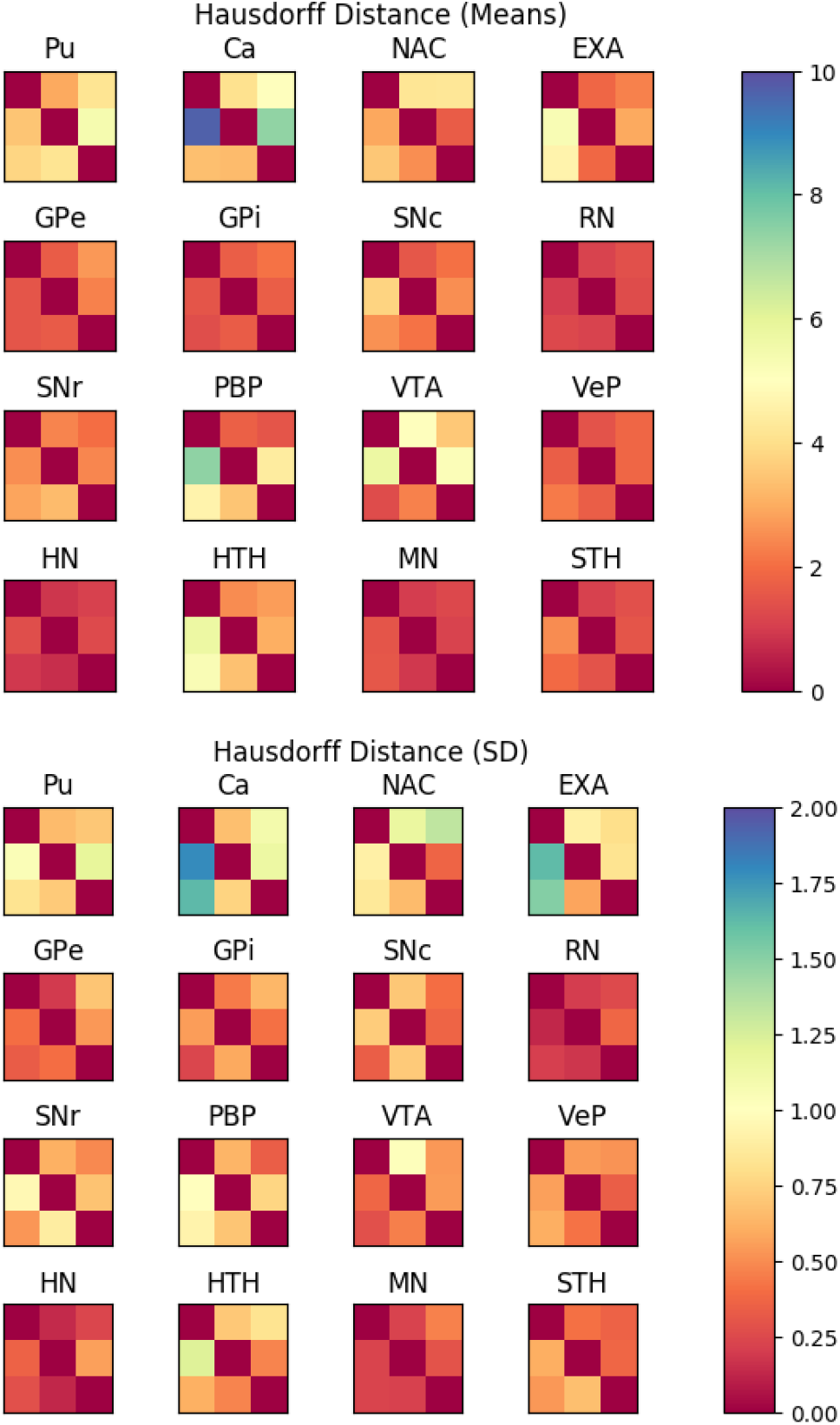
Inter-rater Hausdorff Distances: Means (top) and and standard deviations (SD; bottom) of Hausdorff distances among raters of the 16 main subcortical nuclei included in this atlas. Each row and column correspond to one rater (AN, JMT, and WMP). Each rater labeled subcortical nuclei in each of the eight generated templates. Means and standard deviations were calculated by aggregating Hausdorff distances across templates. Underlying inter-rater Hausdorff coefficients for the different validation templates can be found in the OSF repository (see Data Citations). See Table 1 for acronyms.

**Figure 7:**
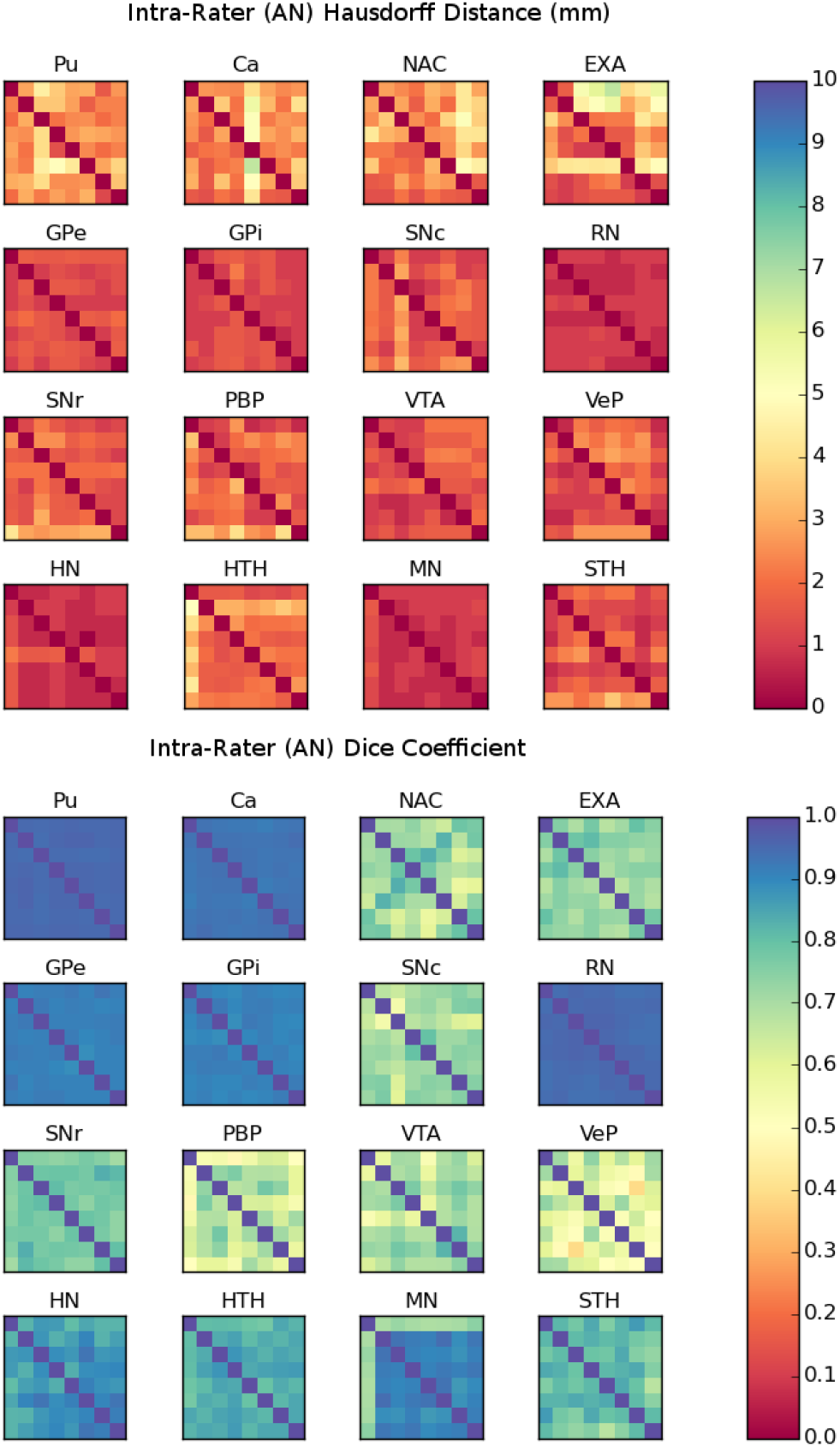
Example Intra-rater Similarities: Intra-rater Hausdorff (top) distance and Dice (bottom) coefficient matrices for each atlas label for observer AN. Each row and column of the matrix represents the results of pairwise comparisons among the eight validation templates. Analogous validation graphs for the other raters can be found in the OSF repository (see Data Citations). See Table 1 for acronyms.

**Table 3:**
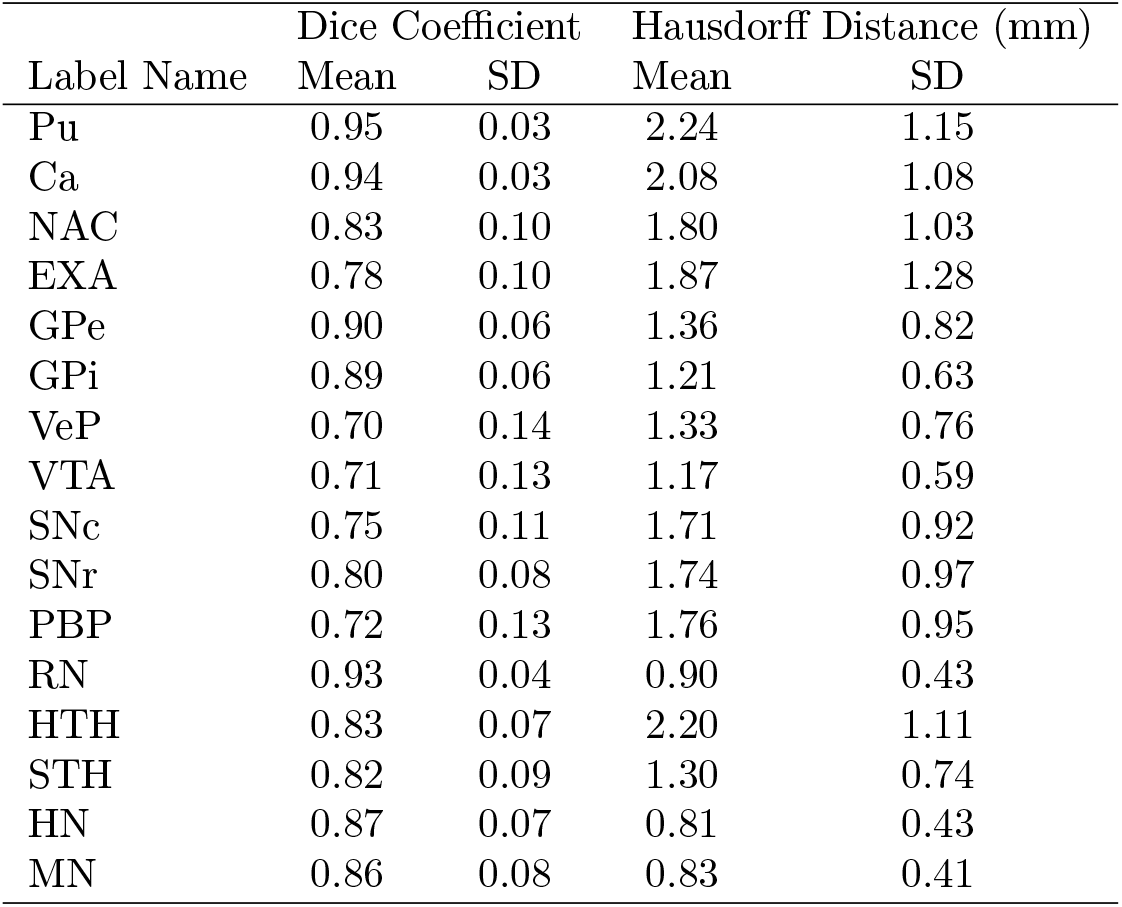
Intra-rater Similarities: Summary statistics for intra-rater Hausdorff distances and Dice coefficients averaged over all observers and templates. See Table 1 for acronyms.

## Acknowledgements

Author contributions: WMP, AN and JMT performed all anatomical region labeling. WMP and JMT wrote the manuscript. We thank Professor Wolfram Schultz (University of Cambridge) for a constructive discussion of the labeling of subcortical nuclei.

This work was funded in part by a Neuroimaging Core of a Conte Center grant from the National Institutes of Health (5P50MH094258-5388). Data were provided in part by the Human Connectome Project, WU-Minn Consortium (Principal Investigators: David Van Essen and Kamil Ugurbil; 1U54MH091657) funded by the 16 NIH Institutes and Centers that support the NIH Blueprint for Neuroscience Research; and by the McDonnell Center for Systems Neuroscience at Washington University.

## References

[1] Christensen, G. E., Rabbitt, R. D. & Miller, M. I. 3d brain mapping using a deformable neuroanatomy. Physics in Medicine & Biology 39, 609 (1994). URL http://stacks.iop.org/0031-9155/39/i=3/a=022.

[2] Thirion, J. P. Image matching as a diffusion process: an analogy with Maxwell’s demons. Medical Image Analysis 2, 243–260 (1998). URL http://www.sciencedirect.com/science/article/pii/S1361841598800224.

[3] Ashburner, J. & Friston, K. J. Nonlinear spatial normalization using basis functions. Human Brain Mapping 7, 254–266 (1999).

[4] Avants, B. B., Duda, J. T., Zhang, H. & Gee, J. C. Multivariate normalization with symmetric diffeomorphisms for multivariate studies. Medical image computing and computer-assisted intervention: MICCAI … International Conference on Medical Image Computing and Computer-Assisted Intervention 10, 359–366 (2007).

[5] Tyszka, J. M. & Pauli, W. M. In vivo delineation of subdivisions of the human amygdaloid complex in a high-resolution group template. Human Brain Mapping 37, 3979–3998 (2016). URL http://onlinelibrary.wiley.com/doi/10.1002/hbm.23289/abstract.

[6] Van Essen, D. C. et al. The WU-Minn Human Connectome Project: An Overview. NeuroImage 80, 62–79 (2013). URL http://www.ncbi.nlm.nih.gov/pmc/articles/PMC3724347/.

[7] Amunts, K. et al. Cytoarchitectonic mapping of the human amygdala, hippocampal region and entorhinal cortex: intersubject variability and probability maps. Anatomy and Embryology 210, 343–352 (2005).

[8] Hawrylycz, M. J. et al. An anatomically comprehensive atlas of the adult human brain transcriptome. Nature 489, 391–399 (2012). URL http://www.ncbi.nlm.nih.gov/pmc/articles/PMC4243026/.

[9] Kamiński, J. et al. Persistently active neurons in human medial frontal and medial temporal lobe support working memory. Nature Neuroscience 20, 590–601 (2017). URL https://www.nature.com/neuro/journal/v20/n4/abs/nn.4509.html.

[10] Wang, S. et al. The human amygdala parametrically encodes the intensity of specific facial emotions and their categorical ambiguity. Nature Communications 8 (2017). URL https://www.ncbi.nlm.nih.gov/pmc/articles/PMC5413952/.

[11] Colas, J. T., Pauli, W. M., Larsen, T., Tyszka, J. M. & O’Doherty, J. P. Distinct prediction errors in mesostriatal circuits of the human brain mediate learning about the values of both states and actions: evidence from high-resolution fMRI. PLOS Computational Biology 13, e1005810 (2017). URL http://journals.plos.org/ploscompbiol/article?id=10.1371/journal.pcbi.1005810.

[12] Haber, S. N. The primate basal ganglia: parallel and integrative networks. Journal of Chemical Neuroanatomy 26, 317–330 (2003). URL http://www.sciencedirect.com/science/article/pii/S0891061803001078.

[13] Hazy, T. E., Frank, M. J. & O’Reilly, R. C. Banishing the homunculus: Making working memory work. Neuroscience 139, 105–118 (2006). URL http://www.sciencedirect.com/science/article/pii/S0306452205005981.

[14] Brown, J., Bullock, D. & Grossberg, S. How the Basal Ganglia Use Parallel Excitatory and Inhibitory Learning Pathways to Selectively Respond to Unexpected Rewarding Cues. Journal of Neuroscience 19, 10502–10511 (1999). URL http://www.jneurosci.org/content/19/23/10502.

[15] Schultz, W., Dayan, P. & Montague, P. R. A Neural Substrate of Prediction and Reward. Science 275, 1593–1599 (1997). URL http://www.sciencemag.org/content/275/5306/1593.

[16] Shen, W., Flajolet, M., Greengard, P. & Surmeier, D. J. Dichotomous Dopaminergic Control of Striatal Synaptic Plasticity. Science 321, 848–851 (2008). URL http://science.sciencemag.org/content/321/5890/848.

[17] Düzel, E. et al. Functional imaging of the human dopaminergic mid-brain. Trends in Neurosciences 32, 321–328 (2009). URL http://www.sciencedirect.com/science/article/pii/S0166223609000757.

[18] O’Doherty, J. P., Cockburn, J. & Pauli, W. M. Learning, Reward, and Decision Making. Annual Review of Psychology 68, 73–100 (2017).

[19] Eshel, N. et al. Arithmetic and local circuitry underlying dopamine prediction errors. Nature 525, 243–246 (2015). URL http://www.nature.com/nature/journal/v525/n7568/full/nature14855.html.

[20] Gerfen, C. R. The neostriatal mosaic: multiple levels of compartmental organization. Trends in Neurosciences 15, 133–139 (1992).

[21] Frank, M. J., Seeberger, L. C. & O’Reilly, R. C. By carrot or by stick: cognitive reinforcement learning in parkinsonism. Science (New York, N.Y.) 306, 1940–1943 (2004).

[22] O’Reilly, R. C. Biologically Based Computational Models of High-Level Cognition. Science 314, 91–94 (2006). URL http://www.sciencemag.org/content/314/5796/91.

[23] Eickhoff, S. B. et al. A new SPM toolbox for combining probabilistic cytoarchitectonic maps and functional imaging data. NeuroImage 25, 1325–1335 (2005).

[24] Frazier, J. A. et al. Structural brain magnetic resonance imaging of limbic and thalamic volumes in pediatric bipolar disorder. The American Journal of Psychiatry 162, 1256–1265 (2005).

[25] Desikan, R. S. et al. An automated labeling system for subdividing the human cerebral cortex on MRI scans into gyral based regions of interest. NeuroImage 31, 968–980 (2006).

[26] Glasser, M. F. et al. The minimal preprocessing pipelines for the Human Connectome Project. NeuroImage 80, 105–124 (2013).

[27] Avants, B. et al. Multivariate Analysis of Structural and Diffusion Imaging in Traumatic Brain Injury. Academic Radiology 15, 1360–1375 (2008). URL http://www.sciencedirect.com/science/article/pii/S1076633208003954.

[28] Avants, B. B., Epstein, C. L., Grossman, M. & Gee, J. C. Symmetric Diffeomorphic Image Registration with Cross-Correlation: Evaluating Automated Labeling of Elderly and Neurodegenerative Brain. Medical image analysis 12, 26–41 (2008). URL http://www.ncbi.nlm.nih.gov/pmc/articles/PMC2276735/.

[29] Avants, B. B. et al. The optimal template effect in hippocampus studies of diseased populations. NeuroImage 49, 2457–2466 (2010).

[30] Yushkevich, P. A. et al. User-guided 3d active contour segmentation of anatomical structures: significantly improved efficiency and reliability. NeuroImage 31, 1116–1128 (2006).

[31] Yushkevich, P. A. et al. Quantitative comparison of 21 protocols for labeling hippocampal subfields and parahippocampal subregions in in vivo MRI: towards a harmonized segmentation protocol. NeuroImage 111, 526–541 (2015).

[32] Van Leemput, K. et al. Automated segmentation of hippocampal sub-fields from ultra-high resolution in vivo MRI. Hippocampus 19, 549–557 (2009). URL http://onlinelibrary.wiley.com/doi/10.1002/hipo.20615/abstract.

[33] Haber, S. N., Fudge, J. L. & McFarland, N. R. Striatonigrostriatal Pathways in Primates Form an Ascending Spiral from the Shell to the Dorsolateral Striatum. The Journal of Neuroscience 20, 2369–2382 (2000). URL http://www.jneurosci.org/content/20/6/2369.

[34] Alexander, G. E., DeLong, M. R. & Strick, P. L. Parallel Organization of Functionally Segregated Circuits Linking Basal Ganglia and Cortex. Annual Review Neuroscience 9, 357–381 (1986).

[35] Middleton, F. A. & Strick, P. L. Basal ganglia and cerebellar loops: motor and cognitive circuits. Brain Research Reviews 31, 236–250 (2000). URL http://www.sciencedirect.com/science/article/pii/S0165017399000405.

[36] Levy, R., Friedman, H. R., Davachi, L. & Goldman-Rakic, P. S. Differential Activation of the Caudate Nucleus in Primates Performing Spatial and Nonspatial Working Memory Tasks. Journal of Neuroscience 17, 3870–3882 (1997). URL http://www.jneurosci.org/content/17/10/3870.

[37] Robinson, J. L., Laird, A. R., Glahn, D. C., Lovallo, W. R. & Fox, P. T. Metaanalytic connectivity modeling: Delineating the functional connectivity of the human amygdala. Human Brain Mapping 31, 173–184 (2010). URL http://onlinelibrary.wiley.com/doi/10.1002/hbm.20854/abstract.

[38] Pauli, W. M., O’Reilly, R. C., Yarkoni, T. & Wager, T. D. Regional specialization within the human striatum for diverse psychological functions. Proceedings of the National Academy of Sciences 113, 1907–1912 (2016). URL http://www.pnas.org/content/113/7/1907.

[39] Mink, J. W. The basal ganglia: focused selection and inhibition of competing motor programs. Progress in Neurobiology 50, 381–425 (1996).

[40] Brown, J. W., Bullock, D. & Grossberg, S. How laminar frontal cortex and basal ganglia circuits interact to control planned and reactive saccades. Neural Networks 17, 471–510 (2004). URL http://www.sciencedirect.com/science/article/pii/S0893608003002521.

[41] Ikemoto, S. Dopamine reward circuitry: Two projection systems from the ventral midbrain to the nucleus accumbens–olfactory tubercle complex. Brain Research Reviews 56, 27–78 (2007). URL http://www.sciencedirect.com/science/article/pii/S0165017307000756.

[42] Fudge, J. L. & Haber, S. N. The central nucleus of the amygdala projection to dopamine subpopulations in primates. Neuroscience 97, 479–494 (2000). URL http://www.sciencedirect.com/science/article/pii/S0306452200000920.

[43] Olszewski, J. & Baxter, D. Cytoarchitecture of the human brain stem. Cytoarchitecture of the human brain stem. (1954).

[44] Wickens, J. Basal ganglia: structure and computations. Network: Computation in Neural Systems 8, R77–R109 (1997). URL http://www.tandfonline.com/doi/abs/10.1088/0954-898X_8_4_001.

[45] Neafsey, E. J., Hull, C. D. & Buchwald, N. A. Preparation for movement in the cat. I. Unit activity in the cerebral cortex. Electroencephalography and Clinical Neurophysiology 44, 706–713 (1978). URL http://www.sciencedirect.com/science/article/pii/0013469478902055.

[46] Deniau, J. M. & Chevalier, G. Disinhibition as a basic process in the expression of striatal functions. II. The striato-nigral influence on thalamocortical cells of the ventromedial thalamic nucleus. Brain Research 334, 227–233 (1985). URL http://www.sciencedirect.com/science/article/pii/0006899385902148.

[47] Chevalier, G. & Deniau, J. M. Disinhibition as a basic process in the expression of striatal functions. Trends in Neurosciences 13, 277–280 (1990). URL http://www.sciencedirect.com/science/article/pii/016622369090109N.

[48] Bullock, D. & Grossberg, S. Neural dynamics of planned arm movements: Emergent invariants and speed-accuracy properties during trajectory formation. Psychological Review 95, 49–90 (1988).

[49] Humphries, M. D., Stewart, R. D. & Gurney, K. N. A Physiologically Plausible Model of Action Selection and Oscillatory Activity in the Basal Ganglia. Journal of Neuroscience 26, 12921–12942 (2006). URL http://www.jneurosci.org/content/26/50/12921.

[50] Frank, M. J., Samanta, J., Moustafa, A. A. & Sherman, S. J. Hold Your Horses: Impulsivity, Deep Brain Stimulation, and Medication in Parkinsonism. Science 318, 1309–1312 (2007). URL http://science.sciencemag.org/content/318/5854/1309.

[51] Ono, T., Nakamura, K., Nishijo, H. & Fukuda, M. Hypothalamic neuron involvement in integration of reward, aversion, and cue signals. Journal of Neurophysiology 56, 63–79 (1986). URL http://jn.physiology.org/content/56/1/63.

[52] Hazy, T. E., Frank, M. J. & O’Reilly, R. C. Neural mechanisms of acquired phasic dopamine responses in learning. Neuroscience & Biobehavioral Reviews 34, 701–720 (2010). URL http://www.sciencedirect.com/science/article/pii/S0149763409001857.

[53] Lammel, S. et al. Input-specific control of reward and aversion in the ventral tegmental area. Nature 491, 212–217 (2012). URL http://www.nature.com/nature/journal/v491/n7423/full/nature11527.html.

[54] Matsumoto, M. & Hikosaka, O. Lateral habenula as a source of negative reward signals in dopamine neurons. Nature 447, 1111–1115 (2007). URL http://www.nature.com/nature/journal/v447/n7148/abs/nature05860.html.

[55] Ji, H. & Shepard, P. D. Lateral Habenula Stimulation Inhibits Rat Mid-brain Dopamine Neurons through a GABA_a_ Receptor-Mediated Mechanism. Journal of Neuroscience 27, 6923–6930 (2007). URL http://www.jneurosci.org/content/27/26/6923.

[56] Hong, S. & Hikosaka, O. The globus pallidus sends reward-related signals to the lateral habenula. Neuron 60, 720–729 (2008). URL http://www.ncbi.nlm.nih.gov/pmc/articles/PMC2638585/.

[57] Dice, L. R. Measures of the Amount of Ecologic Association Between Species. Ecology 26, 297–302 (1945). URL http://www.esajournals.org/doi/abs/10.2307/1932409.

[58] Taha, A. A. & Hanbury, A. An Efficient Algorithm for Calculating the Exact Hausdorff Distance. IEEE Transactions on Pattern Analysis and Machine Intelligence 37, 2153–2163 (2015).

[59] Huttenlocher, D. P., Klanderman, G. A. & Rucklidge, W. J. Comparing images using the Hausdorff distance. IEEE Transactions on Pattern Analysis and Machine Intelligence 15, 850–863 (1993).

[60] Eskildsen, S. F. et al. BEaST: brain extraction based on nonlocal segmentation technique. NeuroImage 59, 2362–2373 (2012).

## Data Citations

1. Tyszka, J. Michael, Pauli, Wolfgang M., and Nili, Amanda N. Open Science Framework (OSF) doi:10.17605/OSF.IO/JKZWP (2017)

